# Beyond P-values: A Multi-Metric Framework for Robust Feature Selection and Predictive Modeling

**DOI:** 10.1101/2025.10.05.680380

**Authors:** Raelynn Chen, Attri Ghosh, Jie Hu, Yong Chen, Jason H. Moore, Ruowang Li

**Affiliations:** Department of Computational Biomedicine, Cedars-Sinai Medical Center; Department of Biostatistics, Epidemiology and Informatics, University of Pennsylvania

## Abstract

High-dimensional biomedical datasets routinely contain sparse signals embedded among vast, correlated features, making variable selection central to building models that generalize. Although significance-based selection is widely used across modalities (e.g., imaging, EHR, multi-omics), statistical significance does not guarantee predictive utility, and vice versa. Yet few methods unify inferential and predictive evidence within a single selection framework. We introduce MIXER (Multi-metric Integration for eXplanatory and prEdictive Ranking), a domain-agnostic approach that integrates multiple selection metrics into one consensus model via adaptive weighting that quantifies each criterion’s contribution. Through simulation studies, we demonstrate that different selection metrics identified markedly different feature sets whose overlaps depended on the underlying feature distributions and signal strength. Applied to Alzhemier’s disease in UK Biobank, MIXER outperformed every individual criterion, including statistical significance, and generalized to an external disease-specific cohort, Alzheimer’s Disease Sequencing Project, yielding higher discrimination and stronger risk stratification. The MIXER framwork is also modular and readily extends to other selection criteria and data modalities, providing a practical route to more accurate, interpretable, and transportable predictive models.

## 1 Introduction

In contemporary biomedical research, datasets are increasingly high-dimensional. A single brain MRI scan contains millions of pixels, while imaging-derived phenotypes (IDPs) often include tens of thousands of quantitative features [Alfaro-Almagro et al., 2018, Elliott et al., 2018]. Single-cell RNA sequencing typically captures expression profiles of up to 30,000 genes per cell [Heumos et al., 2023], and genome-wide association studies (GWAS) routinely assess millions of single nucleotide polymorphisms (SNPs) [Visscher et al., 2017, Abdellaoui et al., 2023]. Similarly, large number of clinical features in electronic health records are routinely mined for associations with disease prognosis and progression [Miotto et al., 2016, Shickel et al., 2017, Landi et al., 2020]. A shared characteristic of these datasets is the sparsity of true signals amidst a vast number of non-informative background variables. An essential and widely adopted task in analyzing high-dimensional datasets is to build generalizable models that can be extrapolated to future patients or unseen samples to predict their outcomes. However, direct modeling of high-dimensional data without appropriate preprocessing is not only computationally intensive but can also compromise the robustness of downstream inference. As a result, effective selection of important features is essential to reduce dimensionality, improve model interpretability, and mitigate the risk of overfitting.

The general strategy for variable selection in biomedical datasets is to evaluate the statistical significance of each feature using regression-based approaches. For example, in GWAS, the association between each SNP (feature) and a disease phenotype is assessed using logistic or linear regression. The SNPs are ranked by the resulting p-value with statistical significance commonly defined by a Bonferroni corrected p-value threshold of 5 × 10^−8^ [Pe’er et al., 2008, Fadista et al., 2016]. In parallel, machine learning-based variable importance metrics, such as permutation importance, Shapley values, and Gini impurity, offer alternative strategies that can capture complex interactions among variables [Lundberg and Lee, 2017, Breiman et al., 2017, Fisher et al., 2019, Hooker et al., 2021]. However, these methods are inherently model-dependent, often lack reproducibility across different runs, and require large sample sizes and extensive tuning. Moreover, their interpretability is limited in comparison to classical statistical methods, which hinders their routine use in many biomedical contexts.

Despite the central role of statistical significance in variable selection, the primary objective of many biomedical studies is predictive accuracy, such as in disease classification, prognosis, or survival prediction. However, as demonstrated by Lo et al. [2015], statistically significant variables are not necessarily predictive, and conversely, predictive variables may not reach conventional significance thresholds. This disconnect underscores a critical gap: existing variable selection frameworks often fail to jointly consider statistical and predictive utility. Furthermore, predictive performance can be evaluated using a variety of metrics—including area under the curve (AUC), F1 score, and balanced accuracy, with each capturing different aspects of model performance [Richardson et al., 2024]. Relying on a single predictive metric may introduce biases analogous to those stemming from exclusive dependence on p-values. Addressing this methodological gap is essential for developing variable selection strategies that are both statistically robust and practically effective for prediction in high-dimensional biomedical settings.

In this study, we propose the Multi-metric Integration for eXplanatory and prEdictive Ranking (MIXER) framework to enable multi objective variable selection in high-dimensional biomedical data. MIXER adopts a data-driven strategy to integrate multiple selection criteria, including statistical significance and diverse predictive performance metrics, by dynamically evaluating the relative contribution of each criterion to identifying predictive variables. The framework introduces several key innovations that address limitations in existing methods. First, MIXER provides a unified approach that incorporates predictive metrics alongside statistical significance (e.g., p-values), enabling more comprehensive variable selection. Second, unlike traditional multi-objective approaches such as Pareto optimization [Deb, 2011], which yield a set of models optimized for different criteria, MIXER generates a single consensus model that synthesizes information across all metrics. This is particularly advantageous in real-world applications, where practical constraints often necessitate the deployment of a single, interpretable model. Third, MIXER is inherently flexible, allowing the integration of arbitrary selection criteria, making it broadly applicable across diverse biomedical domains and prediction tasks.

We demonstrate using simulated genetic datasets that statistical significance and predictive performance criteria often identify markedly different sets of important features. Models derived from MIXER-selected variables consistently outperform those based solely on p-value selection. We further applied the MIXER framework to develop a genetic risk prediction model for Alzheimer’s disease (AD) using the UK Biobank (UKBB) [Bycroft et al., 2018] and validated its performance in the independent Alzheimer’s Disease Sequencing Project (ADSP) [Leung et al., 2025]. In both datasets, MIXER selected SNPs yielded superior predictive accuracy compared to those selected by statistical significance alone. Together, these results highlight the limitations of relying solely on p-value-based selection and establish MIXER as a robust, generalizable framework that bridges statistical inference and predictive modeling for high-dimensional biomedical data.

## 2 Result

### 2.1 Overview of the Study

The MIXER algorithm is a flexible, multi-objective framework designed to improve the performance, robustness, and generalizability of predictive models by integrating variable selection across diverse evaluation strategies (Figure 1). Traditionally, variable selection has relied on statistical significance, such as p-values, but significance alone does not necessarily translate into predictive accuracy [Lo et al., 2015]. MIXER addresses this limitation by adopting a two-step selection and integration strategy (Figure 1a). Step 1: MIXER uses a data-driven process to evaluate and quantify the quality of multiple variable selection strategies based on their contributions to predictive performance. These strategies can include commonly used significance-based metrics (e.g., p-values), prediction-based measures, or any user-defined metrics (Figure 1b). Step 2: Using a newly proposed predictive importance matrix (PIM), MIXER incorporates the relative importance of different strategies and integrates variables selected across them into a unified predictive model (Figure 1c). This two-step process enables MIXER to leverage diverse selection strategies, resulting in models that are more accurate, robust, and generalizable. We applied MIXER to two independent datasets: UK Biobank (UKBB) and the Alzheimer’s Disease Sequencing Project (ADSP), to evaluate its effectiveness in real-world biomedical contexts (Figure 1d). We assessed performance in terms of predictive accuracy, risk stratification, and biological interpretability, demonstrating that MIXER-selected variables outperform conventional p-value-based approaches across all metrics (Figure 1e).

**Figure 1:**
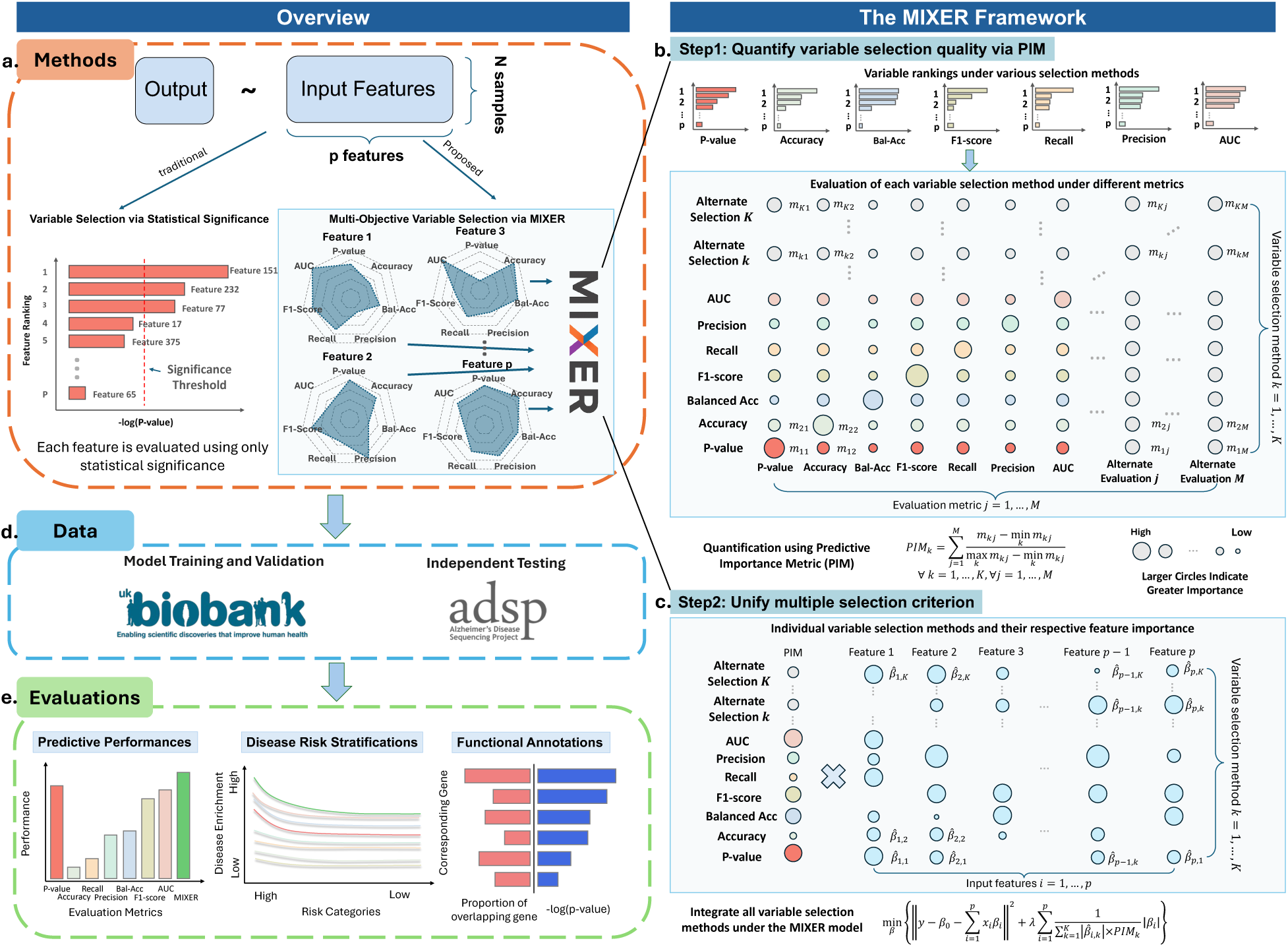
MIXER overview. Panel a illustrates the distinction between the proposed MIXER framework and traditional variable selection methods based on statistical significance. Panels b and c highlight two key elements of the MIXER framework: Panel b depicts the quantification of alternative variable selection criteria, and Panel c demonstrates the integration of multiple selection metrics within the MIXER framework. Panel d summarizes the data structure, including the model training/validation datasets and the external testing dataset. Panel e presents the evaluation components, encompassing predictive performance, risk stratification, and genome-wide functional annotation analyses.

### 2.2 Simulation Study

To evaluate the advantage of MIXER’s multi-objective selection over traditional pvalue-based selection, we conducted a simulation study using simulated genetic datasets with binary outcomes. In each simulation, we generated binary phenotypes using real genetic data from UKBB, and applied both MIXER and a standard p-value thresholding approach to select top-ranking single nucleotide polymorphisms (SNPs). We then trained predictive models using the selected SNPs and evaluated model performance across a diverse set of metrics. To systematically assess performance across feature set sizes, we selected the top 600, 800, and 1000 SNPs based on each method’s ranking. Models trained on MIXER-selected SNPs consistently outperformed those based on p-values alone across all metrics and all selection thresholds (Figure 2). Additionally, alternative selection metrics identify distinct features not captured by p-value–based selection (Supplementary Figure S1). These findings demonstrate that a multi-metric integration approach can lead to more accurate and robust models compare to p-value only selection.

**Figure 2:**
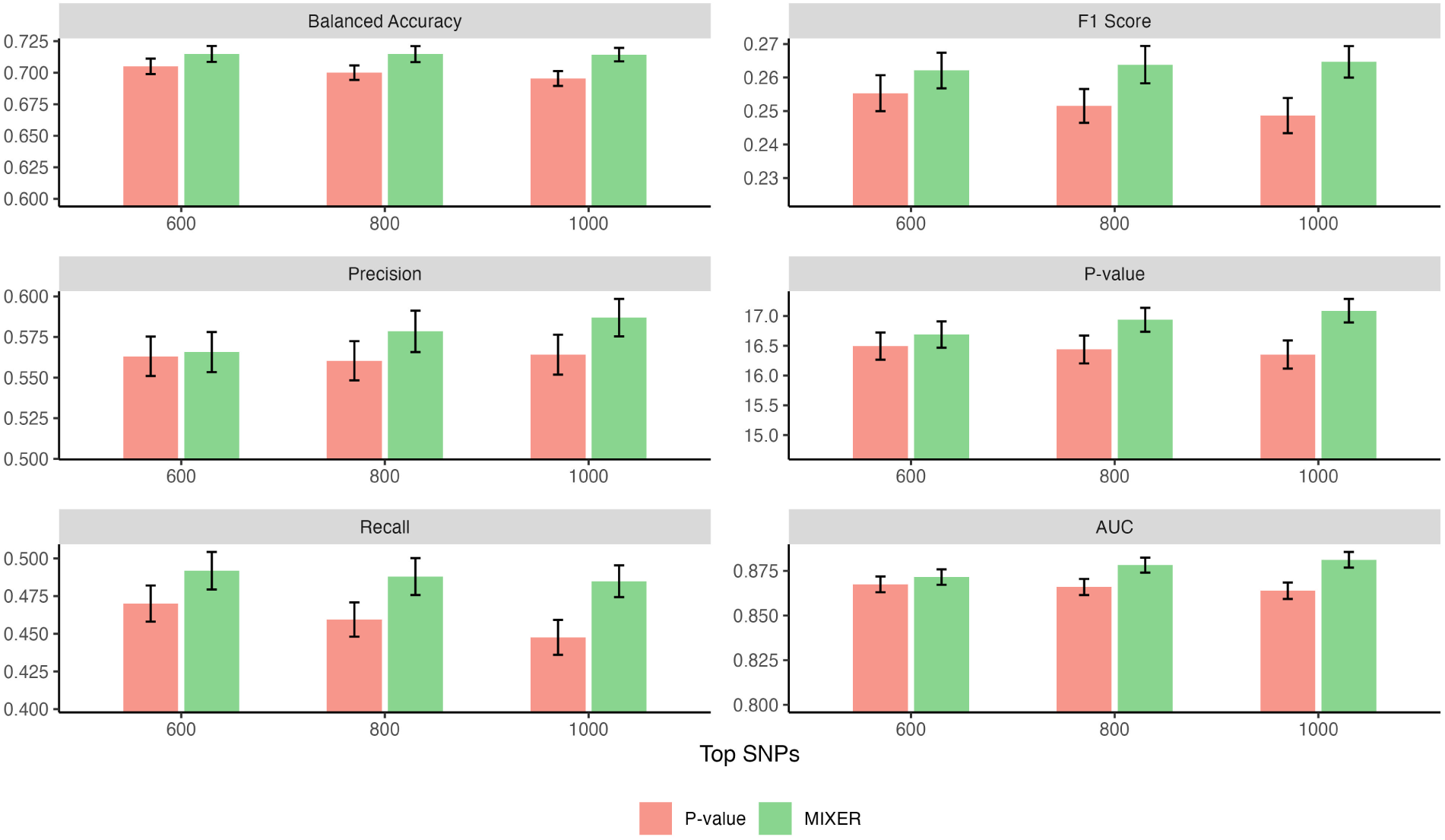
Simulation results comparing p-value-based SNP selection with the MIXER algorithm across six evaluation metrics. For each method, the top 600, 800, and 1000 SNPs were selected and used to train predictive models. Performance was assessed using balanced accuracy, F1 score, AUC, precision, recall, and an overall p-value–based metric.

### 2.3 MIXER Performance Across Cohorts

To assess the predictive performance of MIXER relative to p-value-based and alternative metric-based SNP selection methods, we conducted experiments using the UKBB for training, validation, and internal testing, and the ADSP for external testing.

Figure 3 presents the results across a broad set of evaluation metrics (balanced accuracy, F1 score, precision, p-value, recall, and AUC) for three SNP selection thresholds: top 0.025%, 0.05%, and 0.1%. In the UKBB internal testing results, MIXER consistently outperforms models based on both p-value ranking and any single alternative metric, regardless of the threshold. Importantly, in each case, MIXER either matches or exceeds the best performance achieved by any other method, demonstrating the value of integrating multiple criteria rather than relying on a single selection strategy.

**Figure 3:**
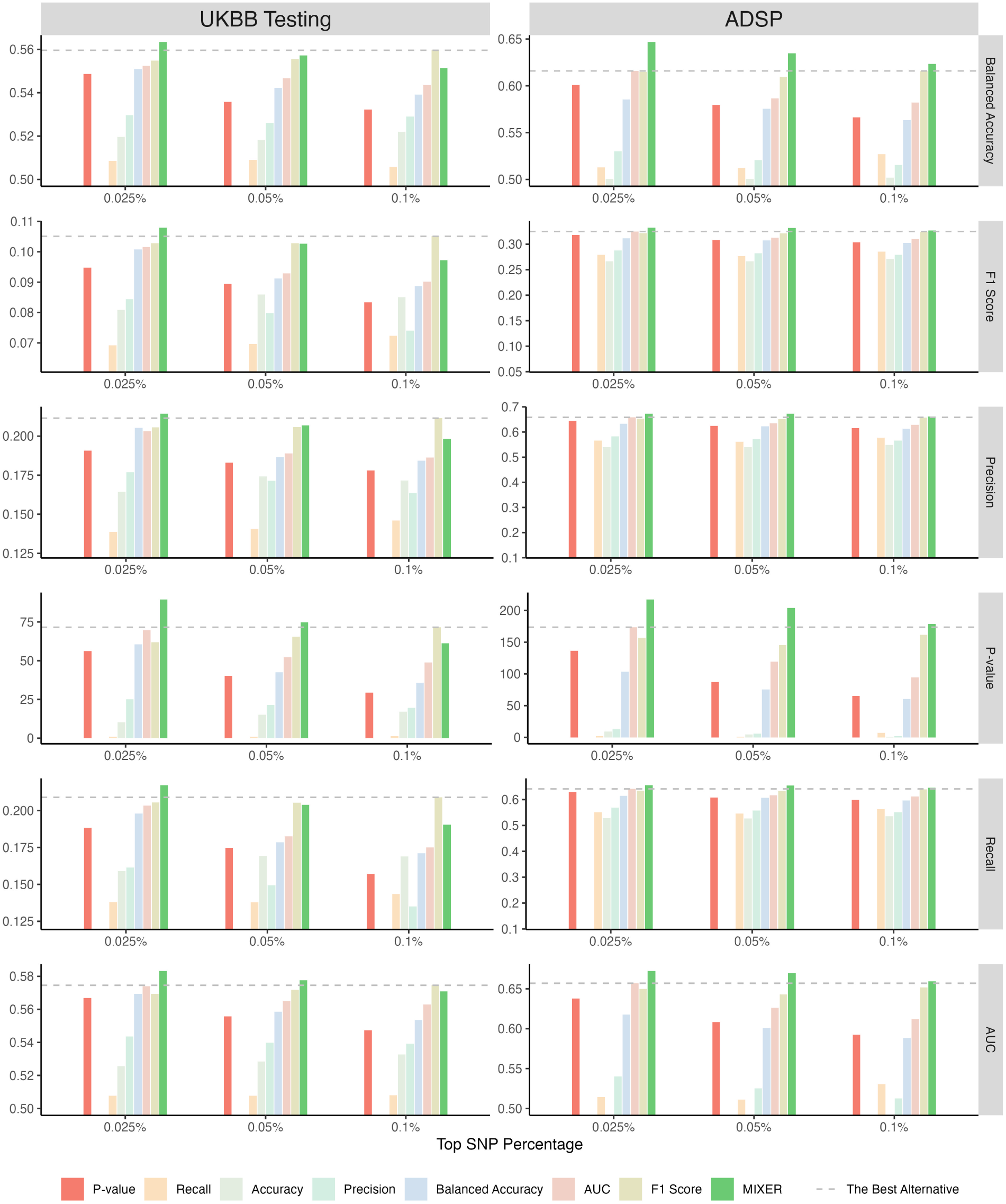
Performance comparison across selection strategies in internal testing in UKBB (left) and external testing in ADSP (right). Each barplot shows model performance using SNPs selected by p-value, MIXER, or one of six predictive metrics (recall, accuracy, precision, balanced accuracy, AUC, F1 score). SNPs were selected based on the top 0.025%, 0.05%, and 0.1% rankings. The gray dashed line indicates the best performance among all individual metrics.

The improvement predictive performance extends to the external ADSP cohort. Despite being trained solely on UKBB data, MIXER models retain improved predictive performance in ADSP. The model consistently outperformed both traditional p-value-based models and the strongest single-metric alternatives.

### 2.4 Characteristics of MIXER-Selected SNPs

To better understand how different selection metrics prioritize SNPs with varying statistical and genetic properties, we analyzed the top 1% of SNPs identified under each criterion. Specifically, we examined the distributions of these SNPs across p-value percentiles, minor allele frequencies (MAF), and effect sizes. The results are summarized in Figure 4.

**Figure 4:**
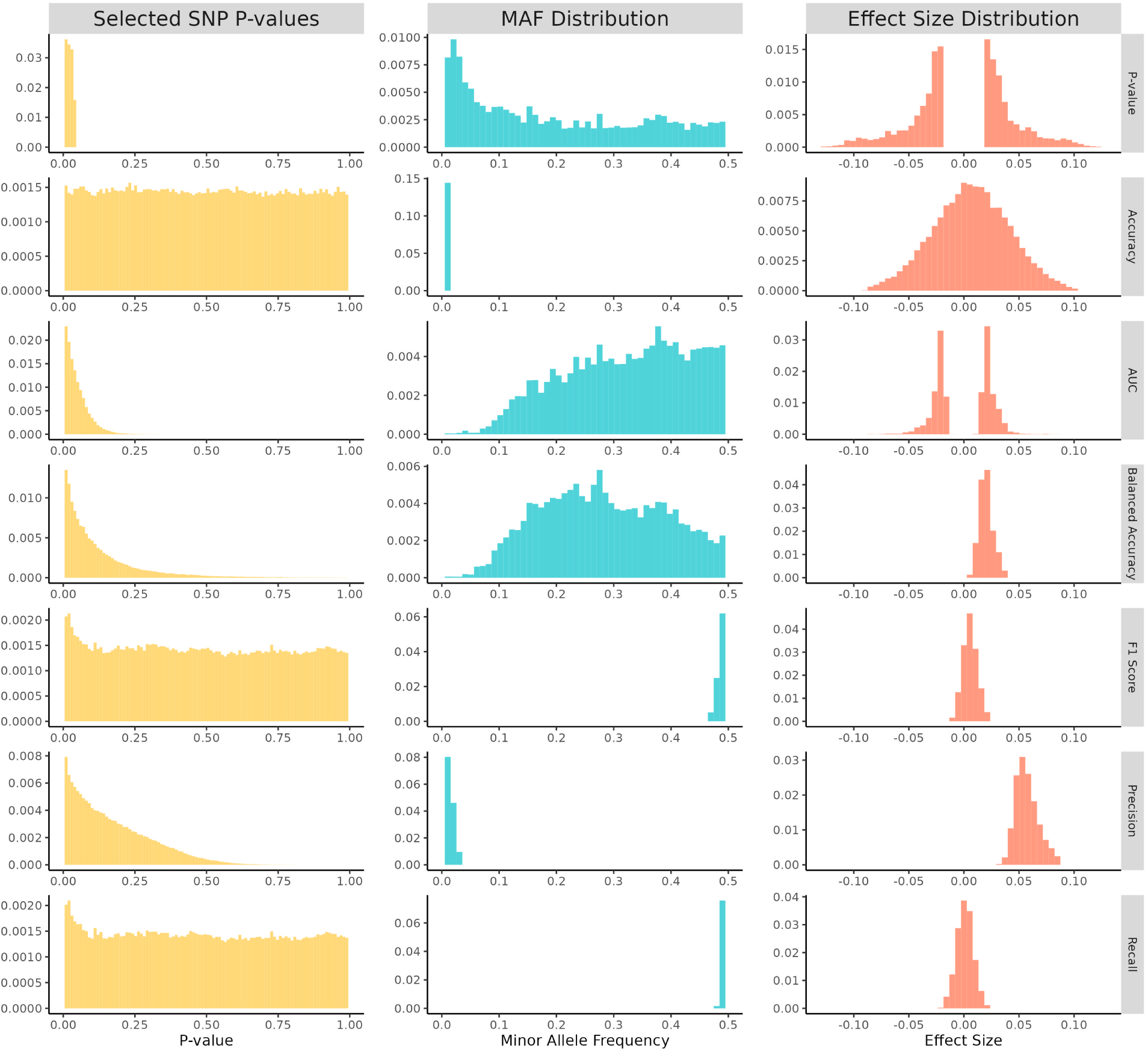
Distributions of the top 1% of SNPs selected under different variable selection metrics. *Left column: P-value distributions of the selected SNPs*. Middle column: Minor allele frequency (MAF) distributions of the selected SNPs. Right column: Effect size distributions of the selected SNPs.

The first column of Figure 4 displays the p-value distributions of the SNPs prioritized by each selection metric. SNPs ranked by p-value show the strongest concentration at values near zero. Among the alternative metrics, AUC exhibits the most pronounced peak at very small p-values, followed by precision and balanced accuracy, which show moderate enrichment near zero. F1 score and recall produce p-value distributions that are close to uniform over [0, 1], but still display a slight peak at zero. Accuracy yields an almost uniform distribution with no visible concentration at small p-values.

The second column shows the minor allele frequency (MAF) distributions of the selected SNPs. SNPs ranked by p-value are enriched at low MAF values (below about 0.1) but still appear across the full range. Accuracy and precision select almost exclusively very rare variants, with mass concentrated near zero. AUC and balanced accuracy display distributions shifted toward common variants, with density increasing toward intermediate and high MAF. F1 score and recall primarily select SNPs with MAF close to 0.5.

The third column shows the effect size distributions of the selected SNPs. For SNPs ranked by p-value and AUC, the distributions are strongly bimodal with modes at moderate positive and negative effects and a clear dip near zero. Accuracy, F1 score, and recall yield approximately bell-shaped distributions: accuracy shows the widest spread and is centered near zero, recall is tightly concentrated around zero, and F1 is bell-shaped but shifted slightly toward positive effects. Balanced accuracy and precision both produce unimodal distributions concentrated on positive effect sizes, with most mass on small to moderate positive effects.

Genome-wide visualization of feature importance from MIXER highlights chromosome 19, which contains the strongest known genetic risk factor for Alzheimer’s disease, as a top-prioritized region (Supplementary Figure S2).

### 2.5 Contribution of Individual Selection Metrics

To better understand the internal weighting within MIXER, we examined the relative contribution of each individual selection metric using the PIM, which quantifies how much each metric influences the final predictive performance of the model.

Figure 5 shows the normalized PIM values across the three SNP selection thresholds. F1 score consistently emerges as the most influential criterion, with its importance growing as more SNPs are included. Balanced accuracy and AUC also show high relative importance across all thresholds. In contrast, recall, accuracy, and precision consistently ranks lowest in importance, while p-value fall in the intermediate range. In summary, these results provide insight into the advantage of MIXER’s adaptive weighting strategy, which emphasizes metrics that contribute most to generalizable predictive performance, rather than assigning equal weight to all metrics by default.

**Figure 5:**
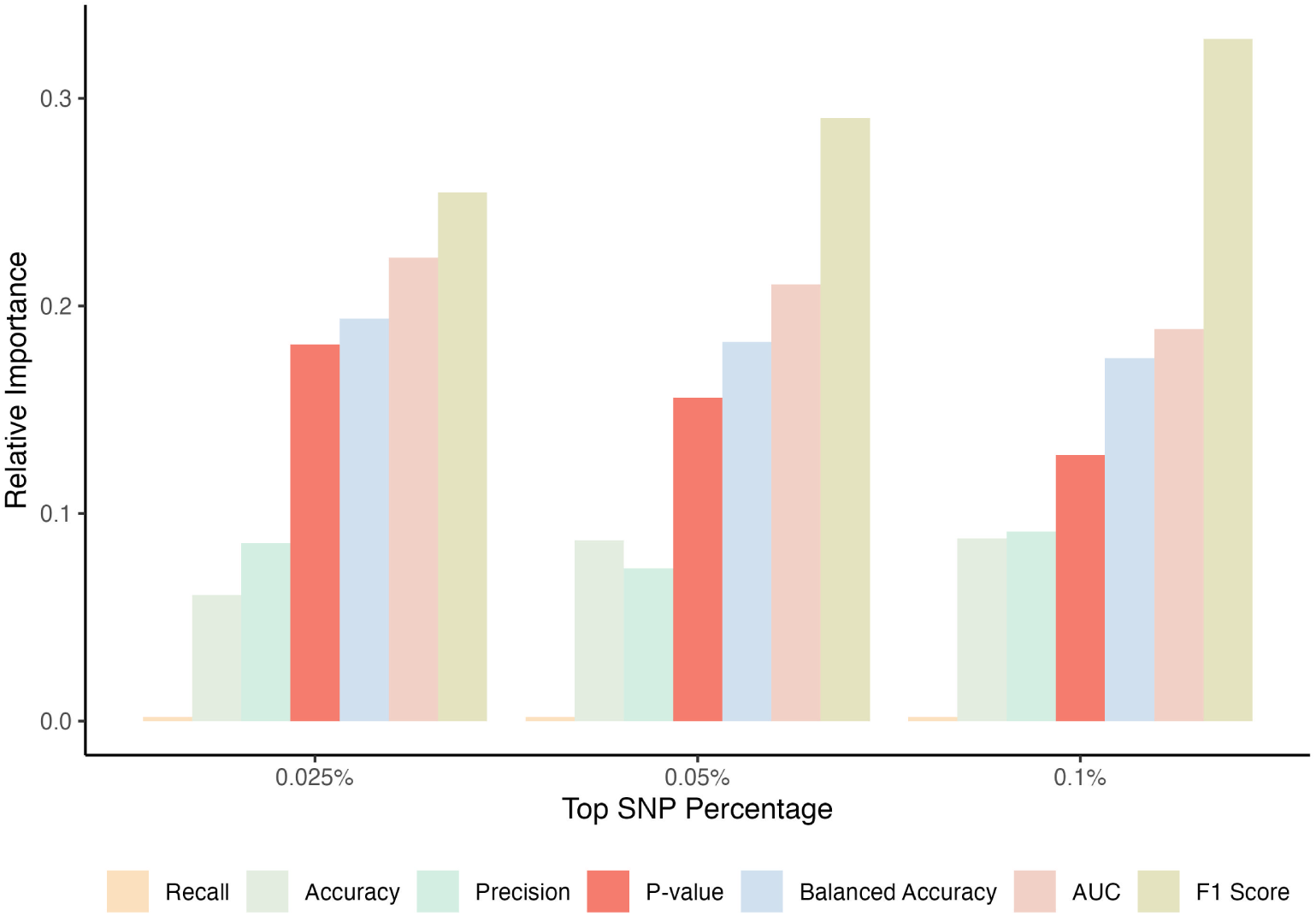
Normalized Predictive Importance Metrics (PIM) across SNP selection thresholds. Bar plots show the relative contribution of each selection metric (p-value, recall, accuracy, precision, balanced accuracy, AUC, F1 score) used in MIXER, based on the top 0.025%, 0.05%, and 0.1% of ranked SNPs. PIM values are normalized to [0,1] for comparison.

### 2.6 Risk stratification Analysis

To assess the clinical utility of MIXER in stratifying individuals by genetic risk, we evaluated odds ratios (ORs) comparing individuals in high-versus low-risk groups. Specifically, we measured the enrichment of individuals in the top 5%, 10%, 15%, and 20% of predicted risk relative to those in the bottom 10%, using models based on SNPs selected by MIXER, p-values, and alternative predictive metrics.

Figure 6 summarizes the risk stratification performance across SNP selection thresholds (top 0.025%, 0.05%, and 0.1%) and datasets (UKBB and ADSP). Across all panels, the ORs consistently decrease as the percentile threshold of the high-risk group increases from 5% to 20%. This trend reflects the expected dilution of risk when expanding the group beyond the most extreme high-risk individuals. Across all conditions, MIXER (green lines) consistently achieves the highest ORs, which indicates better ability to distinguish high-risk individuals. This trend is especially prominent in the ADSP external test set, demonstrating strong generalizability across cohorts.

**Figure 6:**
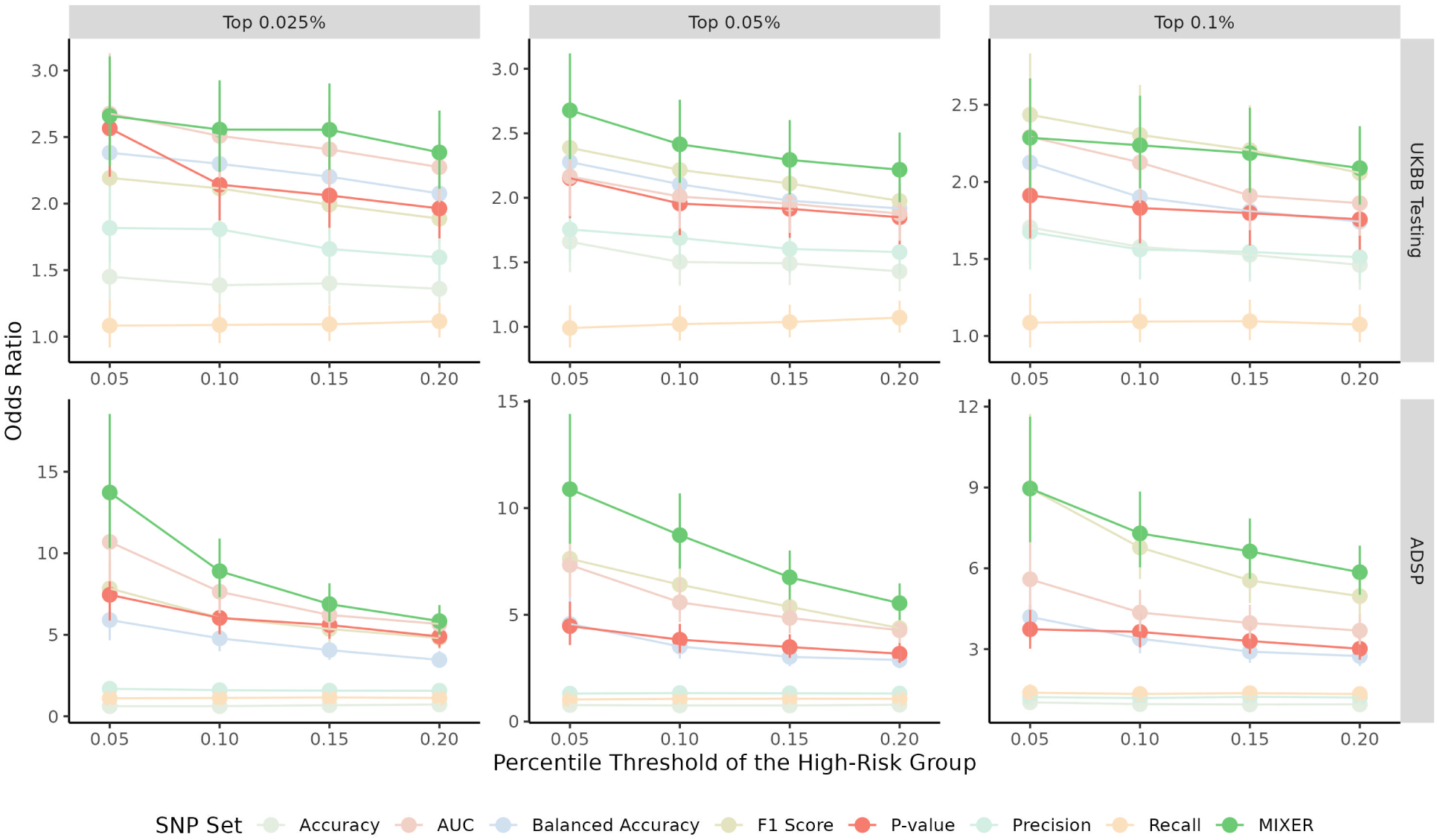
Risk stratification performance across SNP selection thresholds and datasets. Each panel compares the odds ratios (ORs) for individuals in the top 5%, 10%, 15%, and 20% of predicted genetic risk to those in the bottom 10%, based on models trained using SNPs selected via MIXER (green) and alternative methods. The top row shows results from UKBB internal testing; the bottom row shows external validation on ADSP. Columns correspond to three SNP selection thresholds: top 0.025%, 0.05%, and 0.1%. Error bars represent 95% confidence intervals.

SNP sets selected by metrics like AUC, F1 score, and accuracy also outperform p-value-based selection (orange line), which yields relatively flat and lower OR curves. Notably, smaller SNP sets (top 0.025%) yield higher ORs for all settings. This further supports the importance of selection of the most informative variants. The stronger ORs observed in the ADSP dataset show that MIXER captures robust genetic signals that generalize well across datasets.

### 2.7 Functional Annotation Analysis

To interpret the biological relevance of SNPs prioritized by MIXER and alternative selection criteria, we applied functional annotation using the FUMA pipeline Watanabe et al. [2017]. FUMA maps SNPs to genes based on positional and regulatory annotations and performs enrichment analysis to identify trait- and pathway-level associations.

Supplementary Figure S3 summarizes the top traits and phenotypes enriched among SNPs selected using different metrics. Each bar plot highlights the top 10 enriched traits, based on overlapping genes and significance levels. SNPs selected by accuracy and balanced accuracy show strong associations with neurodegenerative and vascular traits, such as cerebrospinal fluid biomarkers (t-tau, p-tau, AB1-42), amyloid pathology, and carotid artery thickness. F1 score, precision, and recall annotated similar features, with emphasis on amyloid burden, cognitive decline, and metabolic processes. P-value and AUC selections are more aligned with structural brain features, including subcortical volume, refractive error, and family history of AD.

Importantly, SNPs selected by MIXER show the strongest enrichment for core Alzheimer’s disease traits, including late-onset AD, APOE status, and age of onset, while exhibiting limited overlap with broader phenotypes.

While all selection metrics identified traits related to Alzheimer’s disease, the specific traits varied across metrics. This further reinforces the importance of using multiple complementary selection criteria.

## 3 Methodology

### 3.1 Overview of the MIXER Methodology

MIXER is a multi-metric variable selection framework that integrates diverse evaluation strategies to identify features with strong predictive utility. It consists of three core stages:

1. **Metric-Specific Feature Selection.** Features are ranked and selected independently based on diverse predictive and statistical criteria (e.g., p-value, AUC, F1 score, etc.).
2. **Quantifying Selection Utility via Predictive Importance Metric (PIM).** The predictive utility of each selection strategy is assessed through standardized modeling and aggregated into a unified importance score.
3. **Adaptive Penalization for Robust Feature Selection.** Selected features are jointly modeled using adaptive LASSO with weights informed by the PIM scores. This promotes both robustness and interpretability, grounded in a strong theoretical framework.

Figure 1 illustrates the overall framework. MIXER adaptively combines the strengths of complementary metrics to improve predictive performance while maintaining interpretability.

### 3.2 Metric-Specific Feature Selection

Let *y* be the outcome of interest and ***x*** = (*x*_1_, · · · *, x_p_*) be the set of candidate features. When *p* is very large, direct joint modeling is computationally and statistically inefficient. As a preprocessing step, MIXER applies univariate models:

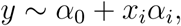

for each *x_i_*, and evaluates its utility using multiple selection criteria. Features are then ranked and the top *q*% are retained per metric. This step mimics real-world practices (e.g., p-value thresholding), but expands beyond p-values to include multiple predictive metrics. As sole reliance on p-values emphasizes statistical association while disregarding predictive accuracy, while reliance exclusively on predictive performance metrics overlooks statistical significance. Multiple independent selection metrics ensures a more balanced tradeoff between statistical association and predictive accuracy.

In this study, we considered a diverse set of metrics, broadly grouped into two categories:

1. Continuous-probability-based metrics:

- Area Under the Receiver Operating Characteristic Curve (AUC): Measures global ranking accuracy.
- P-value: Statistical significance from univariate models.
2. Threshold-dependent classification metrics:

- Accuracy: Overall correctly predicted instances
- Balanced Accuracy: correct predictions while account for class imbalance.
- Recall: Sensitivity to true positives.
- Precision: Positive predictive value.
- F1 Score: Harmonic mean of recall and precision.

For discrete metrics, predictions are binarized using a consistent threshold (e.g., outcome prevalence) throughout our evaluations.

### 3.3 Quantifying Selection Utility via Predictive Importance Metric

To assess which selection strategies yield more predictive features, we introduce the Predictive Importance Metric (PIM). It quantifies the downstream utility of each metric’s selected features as follows:

#### Step 1: Standardization of Feature Sets

1. Let *T_k_* denote the top-ranked features from metric *k*.
2. Let *S_U_* = *T*_1_ ∪ *T*_2_ ∪ · · · ∪ *T_K_* be the union of all selected features, with *p_U_* denote the size of *S_U_*.
3. For each metric *k*, retain the top *p_U_* features to ensure fair comparison.

#### Step 2: Model Evaluation

1. For each standardized feature set *S_k_*, fit the following ridge regression model:

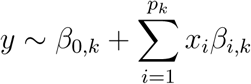

where *y* = (*y*_1_, · · · *, y_N_*), and *x_i_*= (*x*_1*,i*_, · · · *, x_N,i_*). Denote ***β****_k_* = (*β*_0*,k*_*, β*_1*,k*_, · · · *, β_p__k,k_*) as the vector of model parameters, then the loss function can be written as

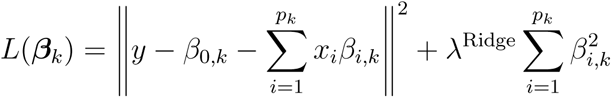
2. Evaluate performance using *M* metrics, e.g., AUC, accuracy, precision, recall, F1 score. Note that the evaluation metrics *M* is not necessarily equal to the *K* selection metrics.
3. Let *m_kj_* denote the performance of the model trained on *S_k_*, evaluated on metric *j*.

#### Step 3. Computing PIM

The PIM is computed as follows:

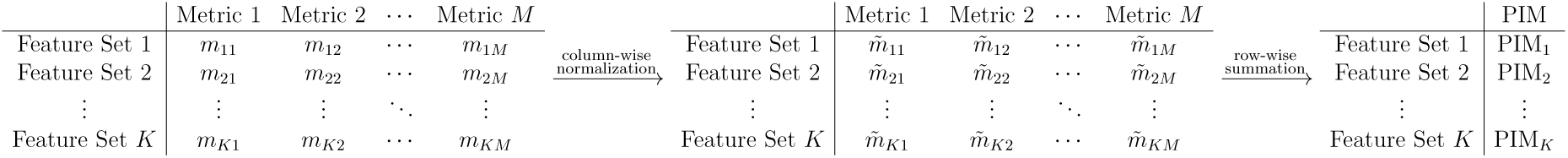

1. Normalize *m_kj_* column-wise to account for scale differences. i.e.,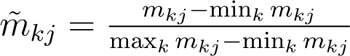.
2. PIM is defined as the row-wise sum of normalized scores: 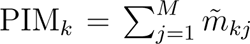.

With higher PIM indicates greater contribution to model performance.

For numerical example of PIM calculation, please refer to Supplementary Table S1-S3.

#### Step 4. Optional threshold calibration (for discrete metrics)

Binary metrics (e.g., F1 score, accuracy) are calibrated by identifying the threshold within the [0, 1] range that yields predicted prevalence closest to the observed prevalence.

In summary, PIM allows MIXER to systematically rank metric-specific strategies by their actual predictive value.

### 3.4 Adaptive Penalization for Robust Feature Selection

To integrate feature-level information across selection strategies, MIXER employs adaptive LASSO, where penalty weights are informed by the PIM-weighted ridge coefficients. Adaptive LASSO was chosen for its ability to improve variable selection and estimation accuracy over traditional LASSO by applying differential penalties that reflect the relative importance of each feature [Zou, 2006].

The adaptive LASSO loss is defined as:

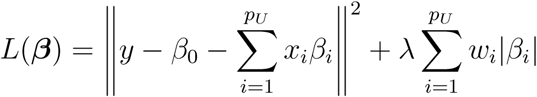

where *p_U_*denotes the number of features in the union set *S_U_*, ***β*** = (*β*_0_*, β*_1_, · · · *, β_pU_*) denotes the regression coefficients, and *w_i_*is the adaptive weight for feature *i*.

The weights *w_i_*are computed as:

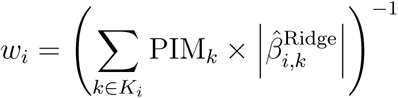

Where *K_i_* = {*k* | 1 ≤ *k* ≤ *K, SNP_i_*∈ *S_k_*} indexes metrics selecting feature *i*, and *β*^^Ridge^ is the ridge estimate of feature *i* in metric *k*, and PIM*_k_* is the predictive importance metric for metric *k*. For numerical example of weights calculation, please refer to Supplementary Figure S2.

This PIM-weighted penalization scheme encourages the inclusion of features that are consistently selected by highly informative strategies (i.e., those with high PIM scores), while applying greater penalties to less informative or weakly supported features. As a result, MIXER adaptively integrates diverse sources of evidence to achieve sparse, robust, and interpretable feature selection.

### 3.5 Simulation Study

We designed our simulation study with two main objectives: (1) to assess whether true causal features could be recovered by multiple feature selection metrics; and (2) to demonstrate that the overlap between features identified by p-values and those selected by alternative metrics is limited, highlighting the potential of alternative metrics to uncover signals missed by traditional approaches. Simulations were conducted using chromosome 22 genotype data from the UK Biobank (UKBB), restricted to individuals of White British ancestry. We excluded SNPs with minor allele frequency (MAF) *<* 0.01, imputation quality *<* 0.8, or genotype missingness *>* 5%, and limited variants to those present in HapMap3. After quality control, 16,470 SNPs remained on chromosome 22. A subset of 5,000 individuals was randomly sampled for simulation.

To generate a binary trait, we specified the log-odds under the linear model:

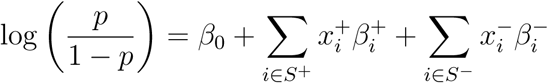

where *S*^+^ and *S*^−^ denote the sets of positively and negatively associated causal SNPs, respectively; *x_i_*^+*/*−^ represents the additive-coded genotype; and *β_i_*^+*/*−^ denotes the corresponding ground-truth effect size. Effect sizes *β*^+*/*−^ were generated in two steps: (1) causal SNPs were randomly selected, with half assigned to the positive group following *β*^+ i.^∼^i.d.^*N* (1, 0.2^2^) and the remainder to the negative group following *β*^−i.^∼^i.d.^*N* (−1, 0.2^2^); and (2) the baseline log-odds *β*_0_ was calibrated to achieve the target prevalence of the binary trait.

We repeated this process 100 times to generate independent replicates. For each replicate:

1. SNPs were ranked using univariate models across multiple selection criteria.
2. For p-value-based selection, a predefined threshold determined the number of SNPs to retain. This number was matched across all other metrics to ensure comparability.
3. MIXER was applied to the selected SNPs from each metric for multivariate selection.
4. MIXER’s performance was compared directly to p-value-based selection under diverse evaluation criteria.

### 3.6 Application of MIXER to UKBB

#### 3.6.1 UK Biobank data (UKBB)

The UK Biobank is a large-scale population-based cohort study that integrates genetic and phenotypic information from approximately 500,000 participants across diverse regions of the United Kingdom [Bycroft et al., 2018]. Genotyping was carried out using either the UK BiLEVE Axiom Array or the UK Biobank Axiom Array, processed in 106 batches, and subsequently imputed with the combined UK10K and 1000 Genomes Phase 3 reference panels. A comprehensive series of quality control (QC) steps was implemented to ensure the reliability of the dataset. At the sample level, we applied the following criteria: (1) inclusion was restricted to individuals with self-reported White British ancestry; (2) individuals were required to have heterozygosity within three standard deviations of the mean; (3) only individuals with self-reported and genetically inferred sex were kept; and (4) only unrelated individuals were retained, with one individual removed from each pair of related participants to minimize bias due to family relationships; relatedness was defined as second-degree or closer, corresponding to an identity-by-descent *π*^ ≥ 0.25. At the SNP level, we excluded variants that met any of the following conditions: (1) duplicated or ambiguous SNPs; (2) imputation INFO score less than 0.8; (3) Hardy–Weinberg equilibrium p-value less than 10^−10^; (4) missingness rate greater than 5%; and (5) minor allele frequency below 1%.

Alzheimer’s disease (AD) status was defined using ICD-10 code G30, extracted from both primary and secondary diagnoses in the UK Biobank (Data-Fields 41202 and 41204). To complement clinical diagnoses and increase power for genetic association studies, we also adopted a proxy-case design as proposed by Liu et al. [2017] and implemented by Marioni et al. [2018]., in which participants with a self-reported parental history of AD or dementia (UKBB Data-Fields 20107 and 20110) were treated as proxy cases, and those with no such family history as proxy controls.

#### 3.6.2 Alzheimer’s Disease Sequencing Project (ADSP)

The Alzheimer’s Disease Sequencing Project (ADSP) is a large-scale, NIH-funded initiative launched in 2012 to sequence and analyze genomes from well-characterized individuals, both with and without Alzheimer’s Disease, in order to uncover genetic risk and protective variants across diverse populations. The multi-phase project includes an initial Discovery Phase, involving whole-genome sequencing (WGS) of approximately 584 family-based samples and whole-exome sequencing (WES) of over 10,000 case-control samples; subsequent Extension and Follow-Up Study phases have expanded sequencing to include tens of thousands of additional genomes from ethnically diverse cohorts. All generated datasets are made available to the broader research community via the NIAGADS DSS repository.

#### 3.6.3 Model training and evaluation

For p-value and AUC metrics, each SNP’s score was directly extracted from the single-SNP regression results. For metrics based on binary predicted labels (e.g., accuracy, F1 score), we applied a sign-informed classification rule: if the regression coefficient (*β*^^^) was positive, individuals carrying at least one minor allele (SNP = 1 or 2) were classified as cases (*y*^ = 1), and major allele homozygotes (SNP = 0) as controls (*y*^ = 0); if *β*^^^ was negative, the assignment was reversed. This approach enabled consistent binary label generation across SNPs for computing discrete classification metrics.

The UK Biobank (UKBB) dataset was randomly split into three subsets: 80% for training, 10% for validation, and 10% for testing. The training set was used for SNP ranking and for fitting ridge and adaptive LASSO models. The validation set was used to compute Predictive Importance Metrics (PIM) and generate adaptive LASSO weights. These two sets were jointly used to estimate the final MIXER model. The testing set was held out to evaluate predictive performance. Additionally, the trained model was independently evaluated in the Alzheimer’s Disease Sequencing Project (ADSP) dataset. For threshold-dependent metrics, the observed disease prevalence was used to guide threshold calibration—13.9% for proxy-AD in UKBB and 54.07% for clinically diagnosed AD in ADSP.

### 3.7 Functional annotations

SNPs prioritized by each selection metric were mapped to genes using positional mapping via the FUMA functional annotation pipeline Watanabe et al. [2017]. For consistency and comparability across metrics, all FUMA parameter settings were held constant throughout the analysis. The resulting gene sets for each metric were then cross-referenced with the NHGRI-EBI GWAS Catalog to assess overlap with previously reported disease-associated loci.

## 4 Discussion

Statistical-significance–based variable selection has long dominated high-dimensional biomedical data analysis because of its simplicity and familiarity, yet significance does not necessarily imply predictive utility. Few studies have systematically (i) tested whether significance translates into out-of-sample performance, (ii) quantified the divergence between significance-and prediction-driven selection, or (iii) proposed a framework that integrates both. Using high-dimensional genetic data as one illustrative use case, we show that significance-based filters often select features that differ markedly from those chosen by prediction-oriented criteria, with the latter yielding more robust generalization. To address this gap, we introduce MIXER, a unified approach that combines significance- and prediction-based variable selections. MIXER outperformed p-value–only pipelines in a population-based biobank and generalized to a disease-specific external cohort, and the framework is readily applicable to other modalities (e.g., EHR, imaging, multi-omics).

Using simulation studies, we found that different selection metrics chose markedly different feature sets and the overlap between p-value selection and prediction-driven selection varied widely across metrics (Figure 4; Supplementary Figure S1). These divergences aligned with differences in minor allele frequency (feature distributions) and effect-size (signal strength). By integrating complementary selection metrics and recovering predictive features that p-value filters may omit, MIXER achieved improved predictive performance across a range of evaluation measures (Figure 2).

Across two distinct cohorts, MIXER outperformed every individual selection criterion (Figure 3). Trained and internally evaluated in a population-based cohort and externally tested in a disease-specific cohort, MIXER’s gains also generalized across different settings, which indicates the model’s robustness to potential cohort shift. Interestingly, the strongest non-MIXER baselines were all prediction-driven, whereas p-value–only selection performed in the middle of the pack. This is confirmed by the analysis quantifying each criterion’s contribution within MIXER ranked p-value fourth of seven (Figure 5). In risk-stratification analyses, which closely align the clinical implementation of a disease prediction model, MIXER produced clearer separation between high- and low-risk groups, while p-value–only models remained intermediate (Figure 6). In summary, MIXER achieves consistent improvements in both predictive performance and actionable risk stratification over significance-only selection. Functional characterization of the features selected by each criterion also revealed complementary, but largely nonoverlapping, biological signals. Although all criteria identified SNPs mapping to Alzheimer’s disease–related traits, their disease-enrichment profiles were distinct. No pair of criteria was enriched for the same set of diseases (Supplementary Figure S3). This pattern indicates that different selection mechanisms prioritize different types of signals, which further supports an integrated strategy, such as MIXER, to aggregate complementary information.

We also acknowledge several limitations that warrant future investigation. First, we evaluated MIXER on genetic datasets across two cohorts. Genetic data was a deliberate choice given its high dimensionality, the availability of large independent cohorts for training and evaluation, and the sparsity of true signals; nevertheless, applications to other modalities, such as EHR, imaging, and multi-omics, are needed to assess generalizability across domains. Second, although this study integrated seven commonly used selection and evaluation metrics, MIXER is also modular. Additional criteria (e.g., PPV, NPV, micro/macro F1, or domain-specific/custom metrics) can be incorporated in future deployments to test their incremental value. Third, MIXER currently employs adaptive lasso for feature/selection integration, which may not capture complex non-linear relationships or higher-order interactions. However, extension to MIXER can incorporate ML-based feature transformations (e.g., interaction expansions, kernels, learned embeddings) or embed non-linear learners precede the framework to enable richer effect modeling [Bengio et al., 2013, Van der Laan et al., 2007].

## Acknowledgment

The ADSP Phenotype Harmonization Consortium (ADSP-PHC) is funded by NIA (U24 AG074855, U01 AG068057 and R01 AG059716).

The ADSP-PHC cohorts include: Adult Changes in Thought (ACT, U01 AG006781, U19 AG066567), the Alzheimer’s Disease Centers (ADC, P30 AG062429 (PI James Brewer, MD, PhD), P30 AG066468 (PI Oscar Lopez, MD), P30 AG062421 (PI Bradley Hyman, MD, PhD), P30 AG066509 (PI Thomas Grabowski, MD), P30 AG066514 (PI Mary Sano, PhD), P30 AG066530 (PI Helena Chui, MD), P30 AG066507 (PI Marilyn Albert, PhD), P30 AG066444 (PI John Morris, MD), P30 AG066518 (PI Jeffrey Kaye, MD), P30 AG066512 (PI Thomas Wisniewski, MD), P30 AG066462 (PI Scott Small, MD), P30 AG072979 (PI David Wolk, MD), P30 AG072972 (PI Charles DeCarli, MD), P30 AG072976 (PI Andrew Saykin, PsyD), P30 AG072975 (PI David Bennett, MD), P30 AG072978 (PI Neil Kowall, MD), P30 AG072977 (PI Robert Vassar, PhD), P30 AG066519 (PI Frank LaFerla, PhD), P30 AG062677 (PI Ronald Petersen, MD, PhD), P30 AG079280 (PI Eric Reiman, MD), P30 AG062422 (PI Gil Rabinovici, MD), P30 AG066511 (PI Allan Levey, MD, PhD), P30 AG072946 (PI Linda Van Eldik, PhD), P30 AG062715 (PI Sanjay Asthana, MD, FRCP), P30 AG072973 (PI Russell Swerdlow, MD), P30 AG066506 (PI Todd Golde, MD, PhD), P30 AG066508 (PI Stephen Strittmatter, MD, PhD), P30 AG066515 (PI Victor Henderson, MD, MS), P30 AG072947 (PI Suzanne Craft, PhD), P30 AG072931 (PI Henry Paulson, MD, PhD), P30 AG066546 (PI Sudha Seshadri, MD), P20 AG068024 (PI Erik Roberson, MD, PhD), P20 AG068053 (PI Justin Miller, PhD), P20 AG068077 (PI Gary Rosenberg, MD), P20 AG068082 (PI Angela Jefferson, PhD), P30 AG072958 (PI Heather Whitson, MD), P30 AG072959 (PI James Leverenz, MD), the Alzheimer’s Disease Neuroimaging Initiative (ADNI), funded by the Alzheimer’s Disease Neuroimaging Initiative (ADNI) (National Institutes of Health Grant U01 AG024904) and DOD ADNI (Department of Defense award number W81XWH-12-2-0012). ADNI is funded by the National Institute on Aging, the National Institute of Biomedical Imaging and Bioengineering, and through generous contributions from the following: AbbVie, Alzheimer’s Association; Alzheimer’s Drug Discovery Foundation; Araclon Biotech; BioClinica, Inc.; Biogen; Bristol-Myers Squibb Company; CereSpir, Inc.; Cogstate; Eisai Inc.; Elan Pharmaceuticals, Inc.; Eli Lilly and Company; EuroImmun; F. Hoffmann-La Roche Ltd and its affiliated company Genentech, Inc.; Fujirebio; GE Healthcare; IXICO Ltd.; Janssen Alzheimer Immunotherapy Research & Development, LLC.; Johnson & Johnson Pharmaceutical Research & Development LLC.; Lumosity; Lundbeck; Merck & Co., Inc.; Meso Scale Diagnostics, LLC.; NeuroRx Research; Neurotrack Technologies; Novartis Pharmaceuticals Corporation; Pfizer Inc.; Piramal Imaging; Servier; Takeda Pharmaceutical Company; and Transition Therapeutics. The Canadian Institutes of Health Research is providing funds to support ADNI clinical sites in Canada. Private sector contributions are facilitated by the Foundation for the National Institutes of Health (https://fnih.org). The grantee organization is the Northern California Institute for Research and Education, and the study is coordinated by the Alzheimer’s Therapeutic Research Institute at the University of Southern California. ADNI data are disseminated by the Laboratory for Neuro Imaging at the University of Southern California, the Memory ang Aging Project at the Knight ADRC (Knight ADRC), supported by NIH grants R01AG064614, R01AG044546, RF1AG053303, RF1AG058501, U01AG058922 and R01AG064877 to Carlos Cruchaga. The recruitment and clinical characterization of research participants at Washington University was supported by NIH grants P30AG066444, P01AG03991, and P01AG026276. Data collection and sharing for this project was supported by NIH grants RF1AG054080, P30AG066462, R01AG064614 and U01AG052410. This work was supported by access to equipment made possible by the Hope Center for Neurological Disorders, the Neurogenomics and Informatics Center (NGI: https://neurogenomics.wustl.edu/) and the Departments of Neurology and Psychiatry at Washington University School of Medicine; the Minority Aging Research Study (MARS, R01 AG22018, R01 AG42210), the National Alzheimer’s Coordinating Center (NACC, U24 AG072122),the National Institute on Aging Late Onset Alzheimer’s Disease Family Study (NIA-LOAD, U24 AG056270), the Religious Orders Study (ROS, P30 AG10161, P30 AG72975, R01 AG15819, R01 AG42210), the RUSH Memory and Aging Project (MAP, R01 AG017917, R01 AG42210), and the National Institute on Aging Genetics of Alzheimer’s Disease Data Storage Site (NIAGADS, U24AG041689) at the University of Pennsylvania, funded by NIA.

The genetic and phenotype datasets generated by UK Biobank analysed are available via the UK Biobank data access process (see http://www.ukbiobank.ac.uk/register-apply/). And our sincere thanks to the participants and researchers of the UK Biobank, who made this effort possible. This project completed under UK Biobank application 86494. We also acknowledge NIH U01 AG066833 for support our study.

## 5 Code Availability

Source code for the MIXER method is freely available for academic purposes at GitHub (https://github.com/Cedars-CIG/MIXER)

