## Supplementary for "Beyond P-values: A Multi-Metric Framework for Robust Feature Selection and Predictive Modeling"

### Supplementary Materials

#### 1 Overlap of Recovered True Signals Across Variable Selection Metrics

To evaluate the overlap and consistency of different variable selection metrics, we conducted a simulation study with known ground truth signals.

In each of the 100 independent simulations, a set of explanatory variables was generated, including a subset of true causal signals and additional noise variables. Each metric (e.g., p-value, accuracy, balanced accuracy, F1 score, precision, recall, AUC) was applied to rank all variables. For each metric, the top $k \in \{600, 800, 1000\}$ranked variables were selected as the "identified signals."

The recovered true signals were then defined as the intersection between the selected variables and the true signal set. For each metric pair, we computed the overlap of recovered true signals, and summarized these results as pairwise Venn diagrams. Each Venn diagram represents the average overlap proportions across 100 simulations.

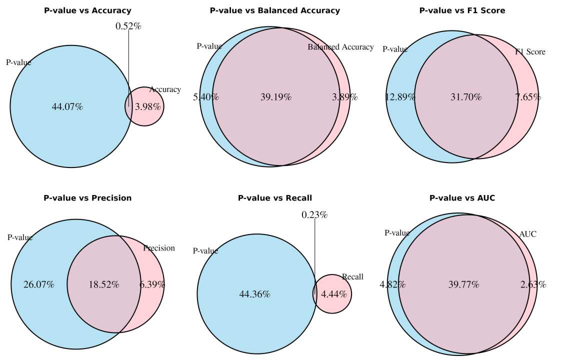


(a) Top 600 set identified signal

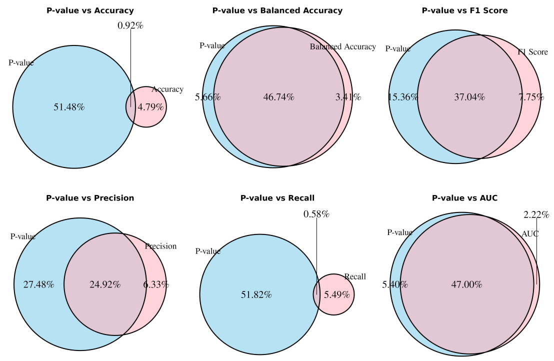


(b) Top 800 set identified signal

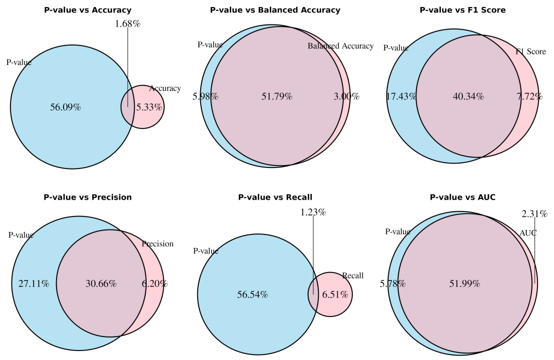


(c) Top 1000 set identified signal

Figure S1: **Average pairwise overlaps of recovered true signals across 100 simulations.** *Each panel corresponds to a different selection threshold: (a) top 600, (b) top 800, and (c) top 1000 variables ranked by each metric. For each metric pair, the Venn diagrams show the average proportion of true signals uniquely or jointly identified by the two metrics. Blue and pink circles represent the true signals recovered by the first and second metric, respectively.*

### 2 Illustration of Predictive Importance Metric

To illustrate the computation of the Predictive Importance Metric (PIM) in practice, we present a numerical example using real data. For demonstration, we selected the top 1% of SNPs ranked by a collection of variable-selection metrics. Table S1 reports the raw metric values for these SNPs. Table S2 shows the result after column-wise normalization, where each metric is rescaled relative to its minimum and maximum across feature sets. Finally, Table S3 provides the resulting PIM scores obtained through row-wise summation of the normalized metrics. Together, these tables provide a step-by-step illustration of the transformation from the original metric matrix to the final PIM values.

| | **Accuracy** | **Bal-Acc** | **F1 Score** | **Precision** | **P-value** | **Recall** | **AUC** |
|---|---|---|---|---|---|---|---|
| **Accuracy** | 0.7706 | 0.5185 | 0.0852 | 0.1727 | 13.3016 | 0.1682 | 0.5263 |
| **Bal-Acc** | 0.7787 | 0.5216 | 0.0862 | 0.1810 | 28.6200 | 0.1645 | 0.5472 |
| **F1 Score** | 0.7798 | 0.5395 | 0.1037 | 0.2092 | 69.6973 | 0.2057 | 0.5717 |
| **Precision** | 0.7796 | 0.5131 | 0.0769 | 0.1663 | 11.1134 | 0.1430 | 0.5295 |
| **P-value** | 0.7795 | 0.5171 | 0.0812 | 0.1735 | 19.1735 | 0.1527 | 0.5379 |
| **Recall** | 0.7671 | 0.5052 | 0.0726 | 0.1494 | 3.1872 | 0.1413 | 0.5142 |
| **AUC** | 0.7809 | 0.5234 | 0.0874 | 0.1850 | 30.3032 | 0.1656 | 0.5479 |

Table S1: **Original metric matrix for the top 1% SNPs.** *Each entry represents the raw value of a selection metric (columns) for a feature set (rows) before any normalization.*

| | **Accuracy** | **Bal-Acc** | **F1 Score** | **Precision** | **P-value** | **Recall** | **AUC** |
|---|---|---|---|---|---|---|---|
| **Accuracy** | 0.2549 | 0.3873 | 0.4044 | 0.3888 | 0.1521 | 0.4174 | 0.2112 |
| **Bal-Acc** | 0.8412 | 0.4796 | 0.4360 | 0.5285 | 0.3824 | 0.3604 | 0.5751 |
| **F1 Score** | 0.9196 | 1.0000 | 1.0000 | 1.0000 | 1.0000 | 1.0000 | 1.0000 |
| **Precision** | 0.9000 | 0.2317 | 0.1373 | 0.2822 | 0.1192 | 0.0270 | 0.2661 |
| **P-value** | 0.8941 | 0.3482 | 0.2766 | 0.4032 | 0.2404 | 0.1772 | 0.4127 |
| **Recall** | 0.0000 | 0.0000 | 0.0000 | 0.0000 | 0.0000 | 0.0000 | 0.0000 |
| **AUC** | 1.0000 | 0.5309 | 0.4752 | 0.5954 | 0.4077 | 0.3784 | 0.5870 |

Table S2: **Column-wise normalized metric matrix for the top 1% SNPs.** *Each column has been rescaled to the* $[0, 1]$ *range to place all metrics on a comparable scale prior to PIM computation.*

| | **PIM** |
|---|---|
| **Accuracy** | 2.2160 |
| **Bal-Acc** | 3.6032 |
| **F1 Score** | 6.9196 |
| **Precision** | 1.9635 |
| **P-value** | 2.7523 |
| **Recall** | 0.0000 |
| **AUC** | 3.9746 |

Table S3: **PIM values for the top 1% SNPs.** *The PIM for each feature set is obtained by summing the corresponding normalized metrics across all columns (row-wise summation).*

### 3 Manhattan Plot for feature weights from MIXER model

We visualize inverse of MIXER weights weights $\frac{1}{w_i}$ using a Manhattan-like plot to examine how the MIXER penalty varies across the genome. Since smaller weights correspond to stronger penalization, plotting their reciprocals highlights SNPs that receive less shrinkage and thus exert greater influence in the model. This representation provides an intuitive, genome-wide view of which loci are prioritized by the MIXER, allowing comparison of signal enrichment patterns across chromosomes and different SNP inclusion thresholds.

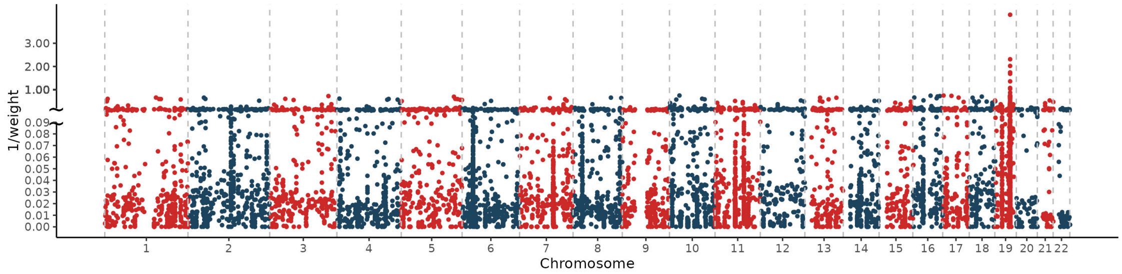


(a) Top 0.025% of SNPs

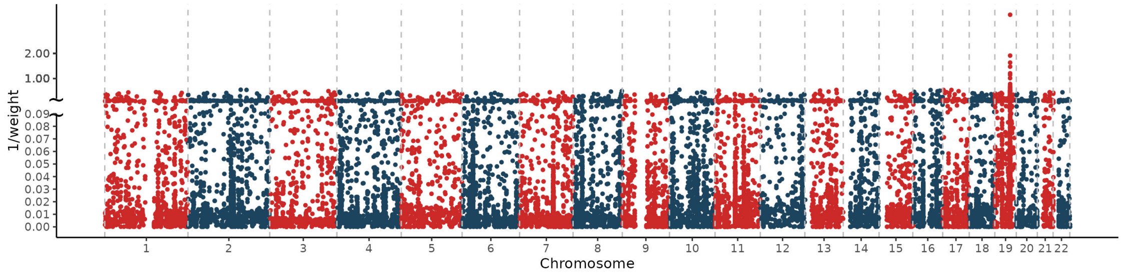


(b) Top 0.025% of SNPs

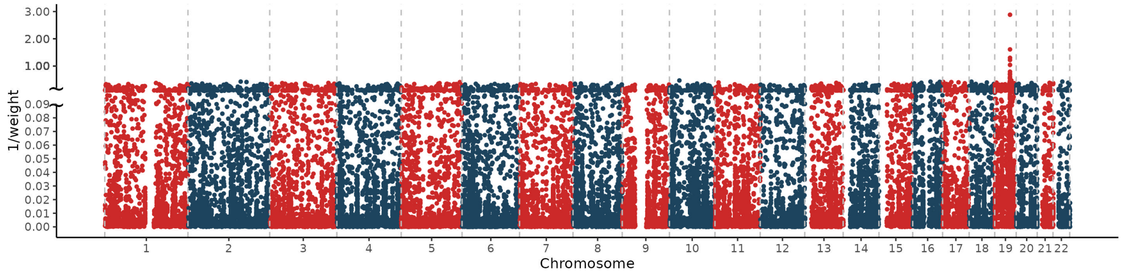


(c) Top 0.025% of SNPs

Figure S2: **Chromosome-wide visualization of the MIXER weights based on SNP subsets of varying selection thresholds.** *Each panel presents the reciprocal of the MIXER weights across chromosomes for (a) the top 0.025%, (b) the top 0.05%, and (c) the top 0.1% SNPs ranked by the initial screening metric. Points represent individual SNPs, alternating in color by chromosome for clarity. Larger values of 1/weight correspond to smaller adaptive weights, indicating stronger penalization and lower relative importance in the MIXER model.*

### 4 FUMA results

FUMA maps SNPs to genes based on positional and regulatory annotations and performs enrichment analysis to identify trait- and pathway-level associations.

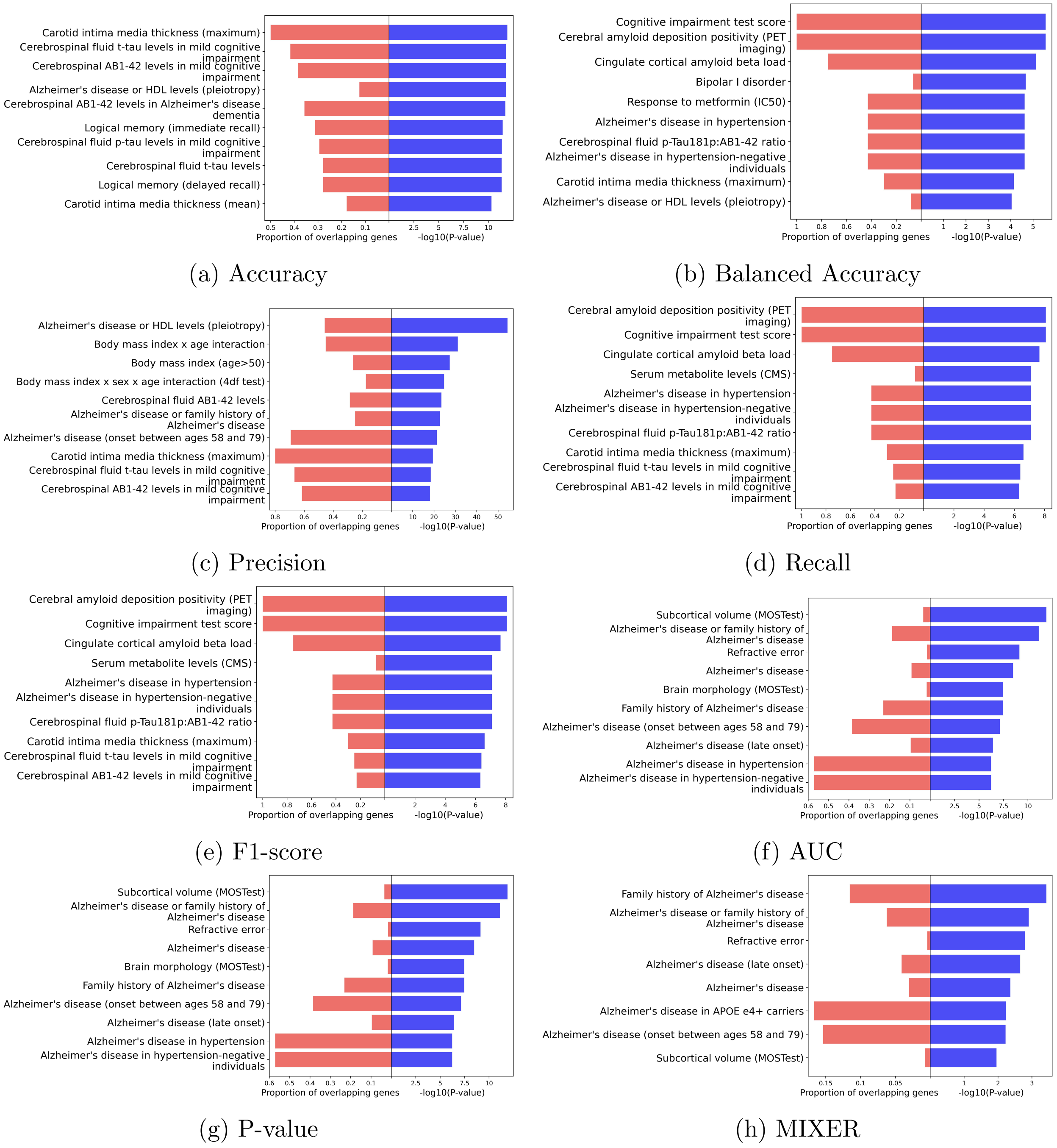


(a) Accuracy (b) Balanced Accuracy (c) Precision (d) Recall (e) F1-score (f) AUC (g) P-value (h) MIXER

Figure S3: **Functional annotation of SNPs selected by different criteria using FUMA.** *Each panel presents enrichment results for SNP sets selected by a given metric. Bars indicate the proportion of overlapping genes associated with each trait (left, red) and the statistical significance of enrichment (right,* $-\log_{10} p - value$*, blue). Results are shown for SNPs prioritized by Accuracy (a), Balanced Accuracy (b), Precision (c), Recall (d), F1 Score (e), AUC (f), P-value (g), and MIXER (h), based on Alzheimer's disease GWAS. Top 10 traits are shown per metric.*
